# High-throughput PBK modelling for dermal exposure: a pragmatic approach to predict systemic pharmacokinetics

**DOI:** 10.1101/2025.11.12.687995

**Authors:** Zeynep Edizcan, Stephan Schaller, Lars Kuepfer, René Geci

## Abstract

High-throughput physiologically based kinetic (HT-PBK) modelling provides a mechanistic framework for predicting systemic pharmacokinetics (PK) from *in vitro* and *in silico* data, supporting non-animal chemical safety assessment within Next generation risk assessment (NGRA). Here, we applied HT-PBK modelling to dermal exposure, a key route of human contact with chemicals. Using the skin permeation model of the Open Systems Pharmacology Suite, we simulated systemic PK profiles of 52 compounds based solely on physicochemical properties predicted by quantitative structure-activity relationship (QSAR) models and without any compound-specific *in vitro* measurements. We systematically compared different QSAR tools for lipophilicity, solubility, and other parameters to identify optimal model parameterisation strategies. Across all compounds, the best-performing HT-PBK strategy predicted 75% of Cmax and AUC values within a tenfold range of observed human plasma data extracted from published clinical studies. A systematic tendency toward overprediction of systemic PK was observed, likely due to missing study metadata and the default assumption of fully hydrated skin. Prediction errors were larger for dermal than for oral exposure, reflecting the greater complexity and variability of dermal absorption processes. Nevertheless, key exposure metrics were reproduced within acceptable limits. These results demonstrate the feasibility of fully *in silico*, non-animal HT-PBK modelling for dermal absorption and support its use as a pragmatic tool for exposure-driven safety assessment within NGRA frameworks.

## Introduction

Next generation risk assessment (NGRA) aims to assess the safety of chemicals without reliance on animal data (Dent et al. 2021). It is based on the integration of new approach methodologies (NAMs) including *in vitro* and *in silico* methods within an exposure-driven framework. A central principle of NGRA is that hazard or bioactivity information obtained from NAMs must be interpreted in the context of internal exposure, meaning the concentration of a substance that occurs within the human body under relevant conditions of use or unintentional contact (Wetmore et al. 2012). This requires approaches capable of predicting systemic pharmacokinetics (PK) in the absence of *in vivo* PK data (Han et al. 2024).

Dermal exposure represents a major route by which humans come into contact with chemicals in pharmaceutical, occupational, environmental, and consumer product settings (Poet and McDougal 2002). The extent to which a compound penetrates the stratum corneum and reaches dermal blood vessels determines its systemic availability and varies widely between substances (Akomeah et al. 2007). Quantifying systemic uptake is therefore essential for linking external dermal application to potential biological effects, whether desired in the case of therapeutic agents or undesired in the case of toxicants.

Traditionally, dermal absorption has been studied using *in vitro* assays or *in vivo* animal and human studies. However, such studies are financially and labour intensive or raise consent and ethical concerns. This creates a clear need for predictive, non-animal alternatives. In the cosmetics sector, this need is most pressing because European legislation bans animal testing for cosmetic ingredients while still requiring systemic safety assessment (Taylor and Rego Alvarez 2020). Similar trends are now visible in the pharmaceutical and chemical industries, where there is increasing emphasis on reducing animal use, accelerating testing, and improving human relevance in safety assessment.

Physiologically based kinetic (PBK) modelling provides a mechanistic basis for predicting the absorption, distribution, metabolism, and excretion (ADME) of substances. Applied to dermal exposure, PBK models integrate skin physiology, compound-specific properties, and circulation to predict systemic substance concentrations. Therefore, they are critical for bridging external dermal exposure with internal concentrations to integrate hazard and exposure data within NGRA (Paini et al. 2019; Paini et al. 2021). However, conventional PBK model development requires extensive *in vivo* data for parameterisation and validation, which limits its application when these data are scarce or even fully unavailable.

High-throughput PBK (HT-PBK) modelling addresses this challenge by parameterising PBK models exclusively from *in vitro* and *in silico* data, including properties predicted by machine learning (ML) and quantitative structure-activity relationship (QSAR) models. Such an approach has the potential to provide scalable, and animal-free predictions of systemic kinetics across large numbers of compounds. While we have previously explored HT-PBK modelling for intravenous and oral exposure (Geci et al. 2024), its application to dermal administration has not yet been evaluated.

The present study applies HT-PBK modelling to dermal exposure. To this end, we compiled a large dataset of human *in vivo* plasma concentration-time profiles after dermal administration (52 compounds) and systematically evaluated different strategies for parameterising PBK models with *in silico* input data. All simulations were conducted using the mechanistic skin permeation model implemented in the Open Systems Pharmacology (OSP) Suite, which represents the stratum corneum, viable epidermis, and dermis as diffusion-partition layers according to the framework established by Kasting and Miller (2006) and implemented by Dancik et al. (2013). By benchmarking against *in vivo* PK profiles and quantifying both systematic and random errors, we identified which *in silico* tools provide the most suitable physicochemical parameters and assessed the reliability of HT-PBK simulations for predicting systemic plasma concentrations after dermal exposure.

## Methods

### Dermal PK and physicochemical property data retrieval

An initial literature search was performed to identify PK studies in which plasma PK data were measured in healthy adult humans after single dermal administration. Studies were excluded if compounds were applied to wounds, irritated skin, or if the compounds were highly volatile or prodrugs, or if the compounds were applied under repeated dosing. We only included data for parent compounds, and only when compound concentrations were measured in plasma or serum. Concentration-time profiles were manually extracted from the retrieved literature and digitised using WebPlotDigitizer version 5.2 (Rohatgi 2024).

QSAR models were used to predict physicochemical properties of all compounds for which we had retrieved PK data. The minimal input properties required for modelling dermal absorption using the OSP Suite were lipophilicity, solubility, melting point, boiling point, and vapour pressure (Kuepfer et al. 2016). For lipophilicity predictions, we used LogD values predicted by ADMETLab 3.0 (Fu et al. 2024), ADMETPredictor (SimPlus; https://www.simulations-plus.com/software/admetpredictor), and OPERA (Mansouri et al. 2018), and LogP values predicted by ADMETLab 3.0, SimPlus, VEGA (Benfenati et al. 2013), and OPERA. Solubility values were predicted using ADMETLab 3.0, SimPlus, VEGA, and OPERA. For melting point and boiling point predictions were made using OPERA and ADMETLab 3.0. Lastly, for vapour pressure, we retrieved predicted values from OPERA and VEGA. PBK model parameters not unique to dermal absorption, such as clearance and fraction unbound, were parameterised in the same way as reported previously (see “Best *in silico*-free" strategy; Geci et al. 2024) to enable comparison of results to previous reports.

### Software and model simulations

To simulate permeation across and within different layers following dermal administration of compounds, we used the skin permeation model of the OSP Suite (Dancik et al. 2013); https://github.com/Open-Systems-Pharmacology/Skin-permeation-model). The model represents the stratum corneum, viable epidermis, and dermis as diffusion-partition layers governed by Fick’s laws of diffusion and partition equilibria, following the biophysical framework described by Kasting and Miller (2006). This mechanistic skin model formed the basis of all HT-PBK simulations in this study and was coupled with the standard whole-body PBK model implemented in PK-Sim version 11.3 (Willmann et al. 2003) to simulate systemic PK. For automated PBK model simulation, we executed models from R using the ospsuite package version 11.3.529 and R version 4.3.1 (R Core Team 2024).

For each collected PK study, a corresponding PBK model simulation was performed by using predicted physicochemical properties together with study-specific parameters such as the formulation of the vehicle, application area, or dosage. When demographic parameters were not indicated in the literature, we used healthy adult males (30 years old, 73 kg, 176 cm) as default settings. Skin thickness values were obtained from studies in literature where histological sectioning, Raman spectroscopy, or ultrasound imaging were used to quantify stratum corneum, viable epidermis, and dermis thicknesses at different anatomical sites (Y. and K. 2002; Sandby-Møller et al. 2003; Egawa et al. 2007; van Mulder et al. 2017; Oltulu et al. 2018; Jeong et al. 2023). As dermal administration studies often provided no application-related study conditions, we assumed healthy skin conditions with a skin surface temperature of 30 °C and fully hydrated stratum corneum. Lastly, for non-occluded vehicles, we assumed a default wind velocity of 16.8 cm/s.

### Predictive performance analysis

The performance metrics were quantified by comparing the results of PBK model simulations against the observed systemic concentration-time data of the corresponding PK studies. We determined the maximum concentration (Cmax) value of simulation and observed data, as well as the area under the curve (AUC) of systemic PK profiles using the linear up-log down method of the PKNCA package version 0.11.0 (Denney et al. 2015). Finally, we calculated the Mean Absolute Log2 Error of Cmax and AUC values of all dataset substances to determine the precision of different prediction strategies as

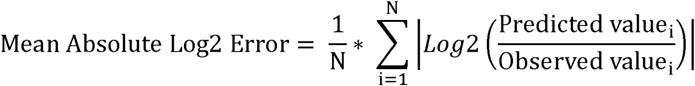

for all N compounds in the dataset. Similarly, we calculated the Median Relative Log2 Error of Cmax and AUC values of all substances to evaluate potential biases of different prediction strategies as

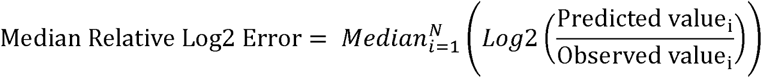

For compounds for which multiple concentration-time profiles had been reported, the median of values across different studies was used to ensure that outliers from individual PK studies did not disproportionately affect the analysis.

## Results

### Retrieval and assessment of dermal PK study data

A total of 64 healthy human PK studies and the *in vivo* plasma concentration-time profiles of 52 unique compounds were collected from the literature (Fig. 1). The identified compounds spanned a broad range of physicochemical properties, including molecular weight, lipophilicity, and solubility (Fig. 1C-E). Most of the dermally applied substances were drugs. The second and third most common groups of compound use were as sunscreen and cosmetics ingredients. Overall, most substance fell into the drug-like chemical space. Application conditions and protocols, such as specific application sites on the body and substance formulations, however, varied strongly between the different clinical studies (Fig. 1F-H).

**Fig. 1.**
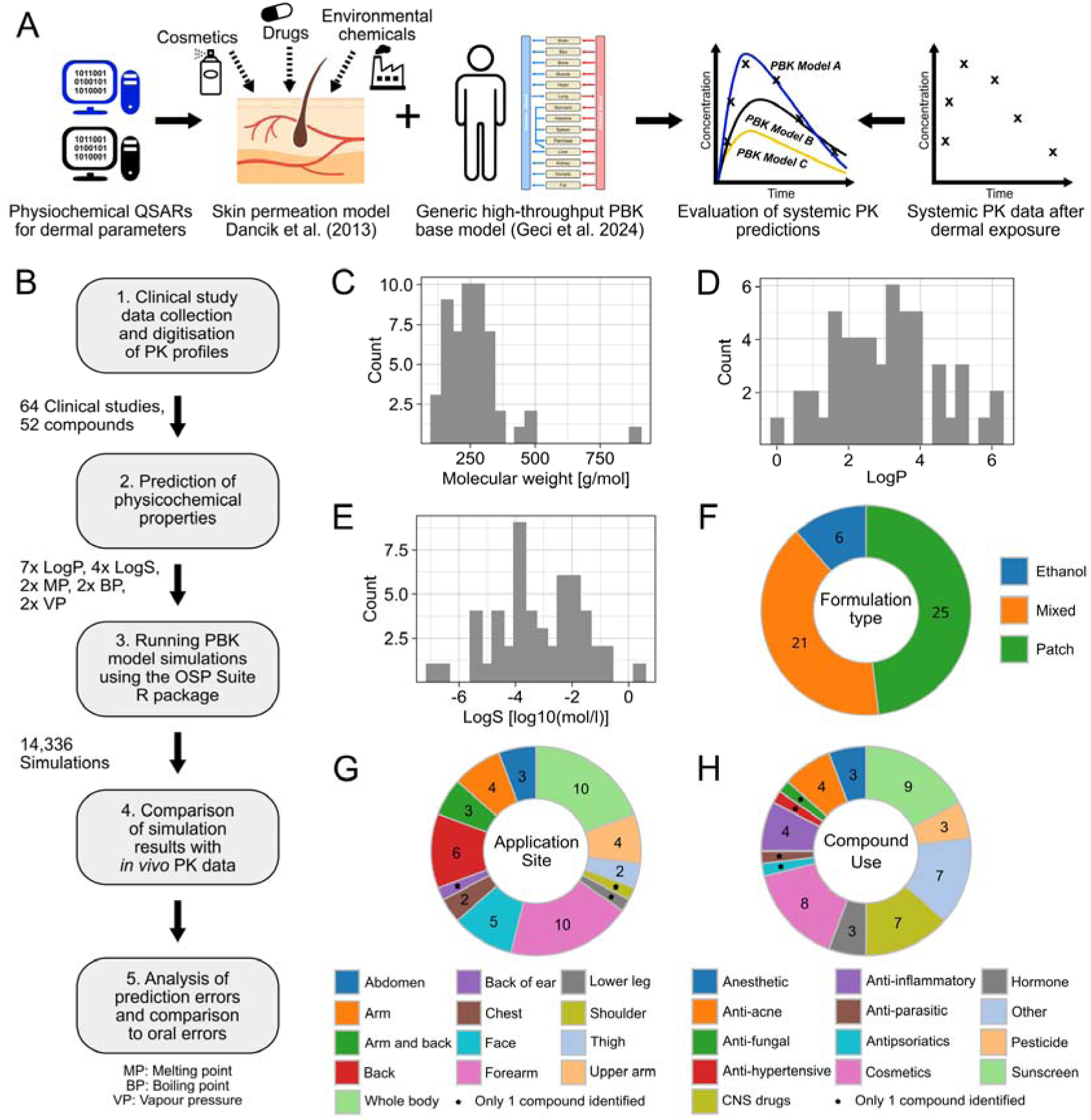
Overview of dermal PK dataset and compound properties. (A, B) Schematic overview of the HT-PBK modelling and analysis workflow for dermally administered compounds. (C-E) Physicochemical properties of the 52 compounds in the collected PK dataset. LogP and LogS values are given as predicted by ADMETLab 3.0. LogS reported as Log10(mol/l). (F-H) The distribution of number of compounds across different study characteristics, such as the type of compound use, formulation type and application site on the body.

For all compounds, we collected the physicochemical property input data required for dermal PBK modelling using different *in silico* prediction tools. Eventually, we obtained lipophilicity predictions from seven different sources, predictions of solubility from four sources, and two predictions for each the melting point, boiling point, and vapour pressure. Since different *in silico* tools use distinct algorithms and training datasets, we combined the various predicted input sources in all possible ways, yielding a total of 224 PBK models per compound. Across 64 clinical studies, this resulted in 14,336 individual PBK model simulations (64 studies × 224 model variants). This systematic evaluation then allowed us to identify which parameterisation strategy provided the most accurate systemic PK predictions. To simplify the evaluation results, we restrict the here shown data to the compound properties which resulted in the largest differences in PK predictions, which were lipophilicity and solubility.

Results for other compound properties, in particular for melting point, boiling point, and vapour pressure, can be found in Supplementary Fig. 1. For those parameters no meaningful differences were observed between the different prediction sources.

### Dermal PBK performance across in silico property predictions

Using the mechanistic skin permeation model of the OSP Suite (Dancik et al. 2013), all dermal absorption simulations were performed by parameterising the model exclusively with *in silico* predicted physicochemical properties. This allowed us to systematically evaluate how different lipophilicity and solubility predictions affected plasma PK simulations. First, we examined the Mean Absolute Cmax and AUC Errors of simulations to evaluate which lipophilicity tools resulted in the most optimal plasma PK predictions after dermal exposure (Fig. 2). With regard to the type of lipophilicity values used, we observed that LogD generally performed worse than LogP. This inferior performance of LogD-based simulations was related to a general bias towards underprediction of the systemic PK data measured *in vivo*. In addition, we also observed differences between the performances of tools predicting the same type of lipophilicity. For instance, LogP values predicted by SimPlus seemed to perform slightly better than values predicted by OPERA.

**Fig. 2.**
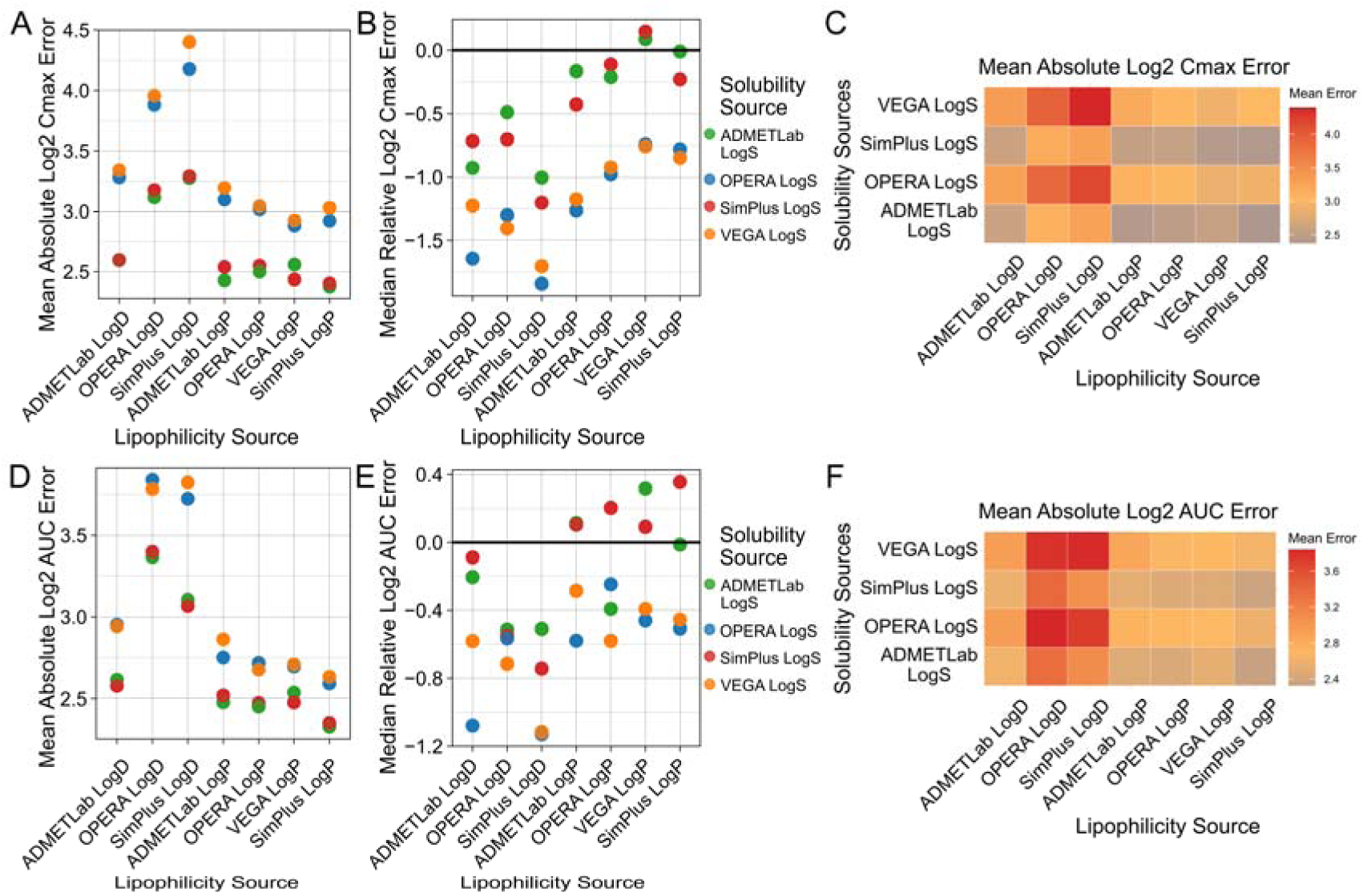
Systemic PK prediction accuracy after dermal exposure across *in silico* input sources. All combinations of physiochemical property prediction tools were used to parameterise the PBK model and evaluated against the collected *in vivo* dermal PK dataset. The first row shows Mean Absolute Log2 Cmax Errors (A), Median Relative Log2 Cmax Errors (B), and a heatmap of Mean Absolute Log2 Cmax Errors (C) for different lipophilicity and solubility prediction sources. The second row shows Mean Absolute Log2 AUC Errors (D), the Median Relative Log2 AUC Errors (E), and a heatmap of Mean Absolute Log2 AUC Errors (F) for different lipophilicity and solubility prediction sources.

Prediction errors also strongly depended on the source of solubility used in PBK model simulations. Again, LogS from SimPlus performed best, together with ADMETLab, and they were followed by VEGA and OPERA. Overall, differences between LogS prediction sources seemed larger than for lipophilicity prediction tools. Heatmaps of the Mean Absolute Log2 Errors of Cmax and AUC are shown across the various combinations of tools used for parameterisation in Fig. 2CF. The overall lowest errors were observed when combining SimPlus LogP with LogS values from SimPlus or ADMETLab.

The simulated plasma concentration-time profiles of five of the best prediction strategies are shown in Fig. 3 for all compounds of the dermal PK dataset. For some substances, the predicted PK profiles showed a PK pattern that was qualitatively comparable to their respective *in vivo* PK profiles. But for other substances the simulated concentration-time profiles strongly differed from the observed PK data. Both strong underpredictions, as well as overpredictions of the *in vivo* data were observed in certain compounds and also the overall dynamics and shapes of profiles deviated in different ways.

**Fig. 3.**
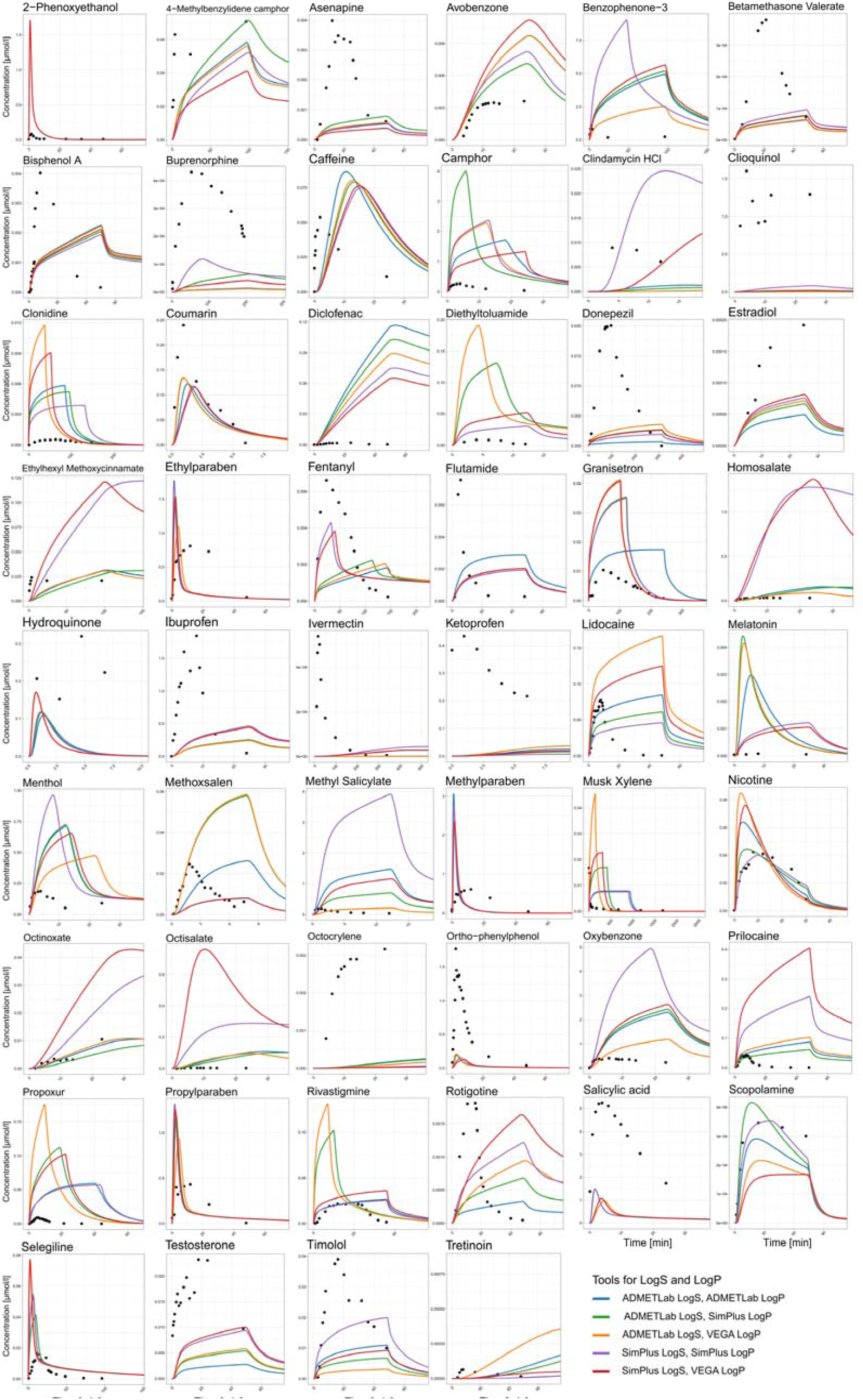
Predicted plasma concentration-time profiles against observed systemic PK data. Shown are the results of five selected best prediction strategies as indicated in the legend for all the compounds in the dataset.

### Correlation between PK prediction error and compound properties

To investigate whether there was a systematic correlation between PK prediction errors and different compounds properties, we further investigated prediction errors the at the individual compound level. However, Cmax and AUC errors showed no consistent association with molecular weight, predicted LogP or LogS, despite wide property ranges (Fig. 4). Only for poorly soluble compounds with low LogS a weak tendency towards underprediction of Cmax and AUC could be observed (Fig. 4CF). However, due to the overall large variability of errors, this trend was not very pronounced. Other predicted properties, such as density, boiling point, melting point, and vapour pressure, showed no clear correlation patterns either (Supplementary Fig. 2). This observation was consistent across different *in silico* tools, indicating that errors were also not related to specific QSAR models.

**Fig. 4.**
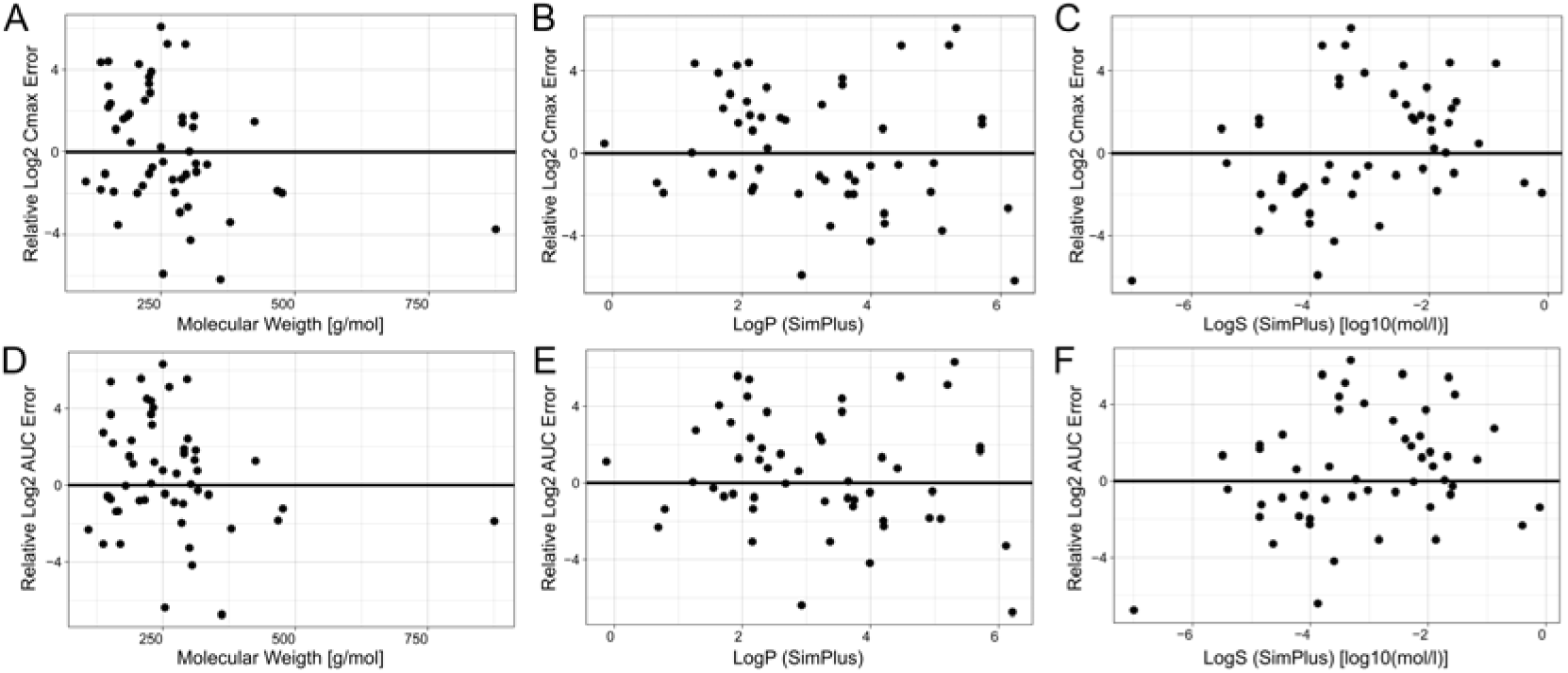
Correlation between prediction error and various compound properties. Results shown are those of the best *in silico* based HT-PBK modelling strategy using SimPlus LogS and LogP. The first row indicates predicted physicochemical parameters including molecular weight, LogP and LogS against Relative Log2 Cmax Errors (A, B, C). The second row shows the same physicochemical parameters against Relative Log2 AUC Errors (D, E, F). Correlations with other compound properties are shown in Supplementary Fig. 2.

### Accuracy of the best systemic PK predictions after dermal application

After determining the best physicochemical property prediction tools for HT-PBK model parameterisation, we eventually assessed in detail how well the best fully *in silico*-based strategy predicted systemic PK after dermal absorption (Fig. 5). We found that 75% of compounds’ Cmax values, as well as 75% of AUC values, were being predicted within the tenfold range. Additionally, we observed that there was a systematic trend towards the overprediction of AUC, although there were also three compounds whose AUC value was underpredicted more than tenfold. This systematic bias was not previously apparent from the Median prediction errors, and we identified that it was caused by the default assumption that the stratum corneum was fully hydrated during chemical exposure. The information whether this was the case or not had not always been provided explicitly in clinical study descriptions, and the assumption of fully hydrated skin conditions was made because hydrated skin is generally more permeable to chemicals, representing a health-protective choice for a pragmatic modelling tool. Since the hydration state of the stratum corneum is a strong determinant of dermal absorption, this assumption likely caused an overprediction of the rate and extent of chemical absorption in HT-PBK simulations. Conversely, restricting the assumption of hydrated skin only to cases explicitly documented in study descriptions, however, would have caused substantial underpredictions of dermal absorption for several compounds (Supplementary Fig. 3).

**Fig. 5.**
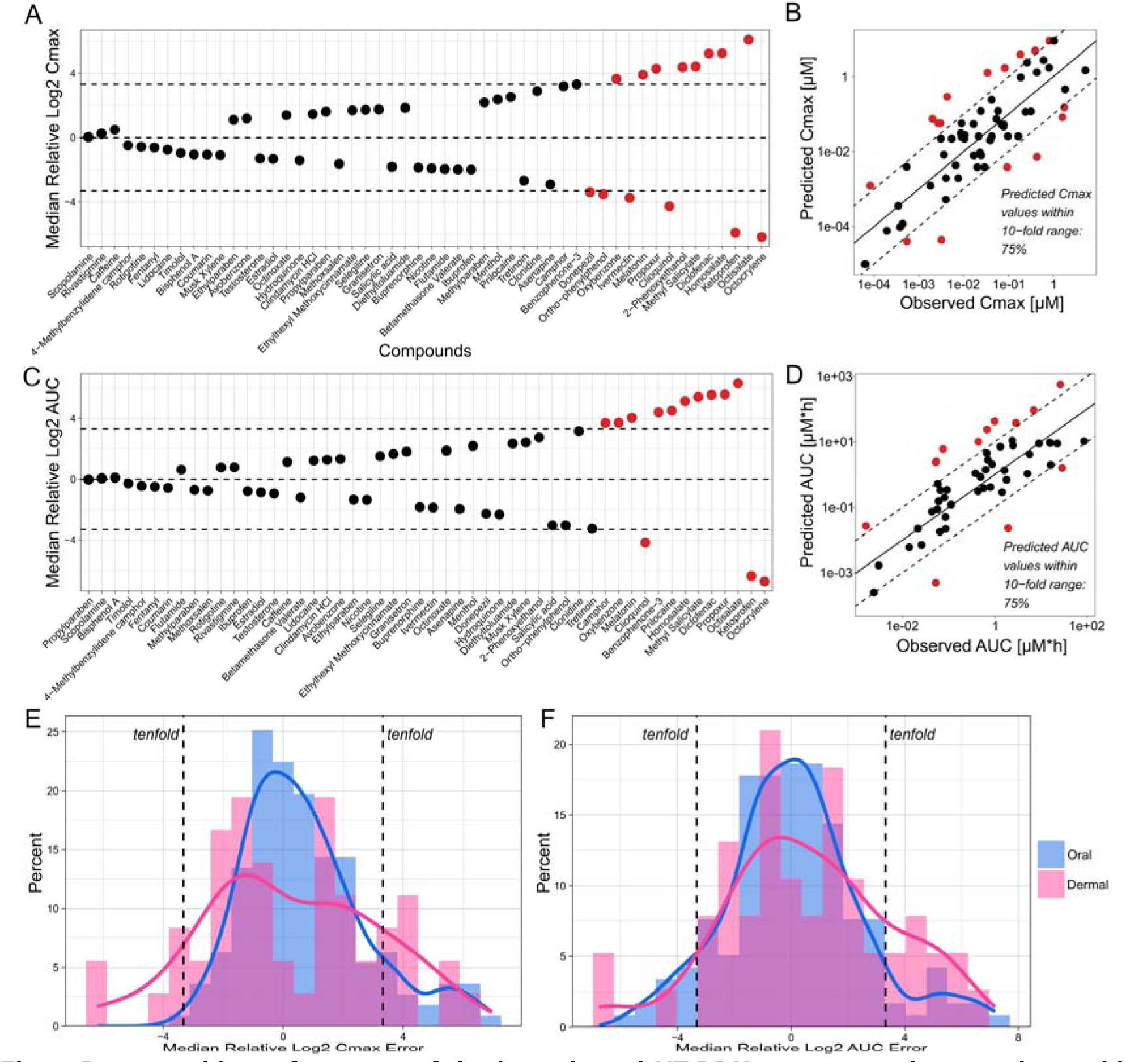
Dataset-wide performance of the best dermal HT-PBK strategy and comparison with oral HT-PBK errors. Model simulations using SimPlus LogS and LogP as the best HT-PBK strategy were evaluated against *in vivo* plasma concentration-time profiles of dermally administered compounds. (A) The Median Log2 predicted/observed Cmax values of each compound in the dataset. (B) Predicted against observed Cmax values of the best strategy for all PK studies. (C) The Median Log2 predicted/observed AUC values of each compound in the dataset. (D) Predicted against observed AUC values of the best strategy for all PK studies. (E) The Median Log2 predicted/observed Cmax values of dermally administered compounds (pink) were compared against errors for prediction of oral administration (blue). (F) The Median Log2 predicted/observed AUC values of dermally administered compounds (pink) were compared against errors for prediction of oral administration (blue). Dashed lines indicate the tenfold error range. Oral HT-PBK prediction errors were taken from Geci et al. 2024.

Finally, we evaluated how the prediction errors of HT-PBK simulations for dermal exposure compared to high-throughput simulations of systemic PK after oral exposure. To this end, we used prediction errors from a previously performed analysis of oral HT-PBK modelling strategies (Geci et al. 2024) and compared the distribution densities of Log2 errors of Cmax and AUC predictions of the two routes of administration against each other (Fig. 5). Despite our efforts to extract a sufficiently large dermal PK dataset, the number of compounds for evaluation after dermal administration was much smaller (54 unique substances) compared to the dataset available for oral administration (161 unique substances). Because of this difference in sample sizes, it proved challenging to compare the distributions of prediction errors of the two exposure routes. Non-normalised Cmax and AUC error distributions are shown in Supplementary Fig. 4. Presumably due to the lower sample size, the variability of dermal prediction errors was relatively large and did not follow a clear normal distribution as it was the case for oral administration. Still, dermal predictions overall showed broader distributions of Cmax and AUC errors, indicating greater variability and uncertainty in systemic PK predictions following dermal exposure (Fig. 5E, F). For comparison, 87% of Cmax and 84% of AUC values were predicted within the tenfold range in our previous analysis (Geci et al. 2024) for oral administration, indicating that systemic PK predictions were generally more accurate for the oral than for the dermal route.

## Discussion

In this work, we used *in silico* predicted physicochemical compound properties and HT-PBK modelling to simulate the systemic PK of 52 dermally administered substances. We assembled a large, curated dataset of healthy human *in vivo* concentration-time data to identify which combination of physicochemical property predictions yield the most suitable parameters for the simulation of systemic PK after dermal absorption, evaluating all possible HT-PBK modelling strategies. Additionally, we examined potential error sources and biases of HT-PBK predictions for compounds relevant to both pharmacological and toxicological applications.

Due to the limited availability of PK studies, less data of dermally administered compounds was retrieved than previously achieved for oral administration. Additionally, we observed that the variability between dermal studies was larger than for intravenous and oral administration. Dermally investigated substances were structurally more diverse, ranging from drugs to sunscreen and cosmetic ingredients, and also the administration procedures differed more substantially for dermal than for oral administration. For instance, across different studies dermal application was performed at different body sites, each with distinct skin-layer thickness and perfusion rates, and using very different vehicles and formulations. In some cases, studies reported that the treated skin had been occluded or hydrated, in other studies no information on such study conditions was reported. Other potentially relevant factors like temperature or wind velocity were also often not reported explicitly. As a result, the large-scale evaluation of HT-PBK model predictions was complicated by additional between-study variability and a lack of study meta data.

One goal of our study was to identify the best compound property prediction tools for high-throughput dermal PBK modelling. Different lipophilicity and solubility sources showed substantial differences in simulation accuracy, whereas there was little variation for other properties such as boiling point, melting point, and vapour pressure. This can be explained by the importance of lipophilicity and solubility for absorption kinetics as both properties are crucial determinants for the permeability of compounds in human skin layers (Cross et al. 2003). The fact that LogP values resulted in better PBK model performance than LogD was expected as the dermal PBK model also formally requests LogP as the lipophilicity input, explaining its superior performance compared to LogD. Nevertheless, we included LogD in this evaluation because it had previously yielded better results for simulation of the whole-body base PBK model. Additionally, we found that there was no clear correlation between the predicted physicochemical property values of substances and their simulation errors for dermal absorption. This observation mirrors the findings of previous work, which also reported the absence of such a relationship when employing experimentally measured physicochemical parameter values (Dancik et al. 2013).

Overall, HT-PBK simulations showed a tendency towards overprediction of systemic PK, particularly regarding AUC values. This bias may partly result from the previously discussed lack of complete clinical study information. For instance, substances might not have been fully released from certain formulations or application vehicles, which could not be accounted for here. Additionally, we made the default assumption that application was performed on fully hydrated skin, which may not have been true for some studies and may have contributed to the here observed overprediction bias. Nonetheless, we proceeded with this assumption to ensure health-protective predictions and to overestimate rather than underestimate *in vivo* substance concentrations. For prospective applications, such factors could, however, be considered more realistically for a given substance and specific exposure scenarios.

Prediction errors of systemic PK seemed generally larger for dermal than for oral exposures, although this comparison was uncertain due to the smaller number of compounds for which dermal PK data was available. In our previous evaluation of oral absorption, 87% of Cmax and 84% of AUC values were predicted within a tenfold range (Geci et al. 2024), whereas the corresponding proportions for dermal exposure were lower (75%). Regarding the congruence between the shape of the predicted PK profiles and of the observed profiles, we also observed much larger differences than for oral administration. Earlier studies indicated that reservoir formation can occur within skin layers, where slow binding or equilibration processes may influence compound release and thereby the absorption kinetics into systemic circulation (Roberts et al. 2004; Anissimov and Roberts 2009). Ignoring such effects might have been a cause for the greater variability observed in dermal absorption compared to the oral absorption of compounds. This supports the interpretation that high-throughput prediction of systemic PK following dermal exposure is more challenging than for oral administration.

Despite this observed misprediction of kinetic profiles, HT-PBK predictions for Cmax and AUC were often within a reasonable range of tenfold within the observed values. Thus, we postulate that while kinetic profiles may not always be predicted accurately, HT-PBK simulations can still provide useful estimates of key exposure metrics relevant for risk assessment. This is plausible, as, for instance, AUC mainly depends on the total absorbed dose and systemic clearance, which may still be predicted correctly even when absorption dynamics are described imperfectly. Ultimately, we conclude that HT-PBK modelling can provide reliable predictions of systemic PK for dermally applied compounds.

All simulations were based on the mechanistic skin permeation model implemented in the OSP Suite (Dancik et al. 2013), which represents diffusion and partition processes across skin layers. Building on this established skin permeation model allowed us to focus on the influence of *in silico* parameterisation rather than on structural model development. In our previous work, we systematically evaluated the same whole-body base PBK model for intravenous and oral administration (Geci et al. 2024). The modular structure of the OSP framework enabled independent validation and optimisation of each exposure route while maintaining consistency within a unified PBK architecture. Building on that foundation, the present analysis could specifically target parameters and processes unique to dermal absorption without re-calibrating systemic components. Such modularity enables integrated simulations across different routes or exposure scenarios, supporting a coherent and extensible framework for quantitative exposure assessment for NGRA.

Several recent studies have developed mechanistic dermal absorption models based on physicochemical properties (Najjar et al. 2021; Clarke et al. 2022; Patel et al. 2022; Dancik et al. 2025; Punt et al. 2025). Other work has addressed uncertainty quantification in dermal PBK modelling, including Bayesian calibration and tiered safety assessment approaches (Li et al. 2022; Hamadeh et al. 2022; Hamadeh et al. 2024). Partition and diffusion coefficients have also been estimated using ML/QSAR and mechanistic methods (Basak et al. 2007; Arora et al. 2022; Hamadeh and Edginton 2023; Narita et al. 2025). However, a systematic, large-scale evaluation of PBK simulations parameterised solely with *in silico* predicted physicochemical properties to predict systemic plasma concentrations after dermal exposure has not been reported.

This study provides such an evaluation, using only *in silico* predicted input parameters to screen systemic PK parameters for 52 dermally applied compounds. In contrast to earlier models focused on a few well-characterised substances, this approach addresses early-stage use cases where compound-specific *in vitro* and formulation data are not available. It establishes a general high-throughput PBK framework applicable across diverse chemical domains, including pharmaceuticals, cosmetics, and industrial chemicals. This supports scalable *in vitro* to *in vivo* extrapolation (Wetmore et al. 2012; Punt et al. 2020) and aligns with current regulatory initiatives on non-animal PBK modelling (OECD 2021; Dent et al. 2021; Ball et al. 2022). It thereby enables quantitative risk assessment without reliance on experimentally measured or animal data, addressing current needs for predictive tools in regulatory and product development contexts.

## Data availability

All data is available in the Supplementary Information.

## Author contributions

Conceptualization: RG; Data curation: ZE, RG; Formal Analysis: ZE, RG; Funding acquisition: SS, LK; Investigation: ZE, RG; Methodology: RG; Software: ZE; Supervision: SS, LK, RG; Visualization: ZE, RG; Writing – original draft: ZE; Writing – review & editing: SS, LK, RG

## Competing interests

Stephan Schaller is founder and managing director of ESQlabs GmbH. All authors declare that they have no conflict of interest.

## Funding

This work was performed in the context of the ONTOX project (https://ontoxproject.eu/) that has received funding from the European Union’s Horizon 2020 Research and Innovation programme under grant agreement No 963845. ONTOX is part of the ASPIS project cluster (https://aspiscluster.eu/).

### Abbreviations

ADME: Absorption, distribution, metabolism, excretion
AUC: Area under the curve
Cmax: Maximum concentration
HT-PBK: High-throughput PBK
LogD: Logarithm of distribution coefficient
LogP: Logarithm of octanol-water partition coefficient
LogS: Logarithm of water solubility
ML: Machine Learning
NAM: New approach methodology
NGRA: Next generation risk assessment
OSP: Open Systems Pharmacology
PBK: Physiologically based kinetic
PK: Pharmacokinetics
QSAR: Quantitative structure-activity relationship

---

**Supplementary Fig. 1.**
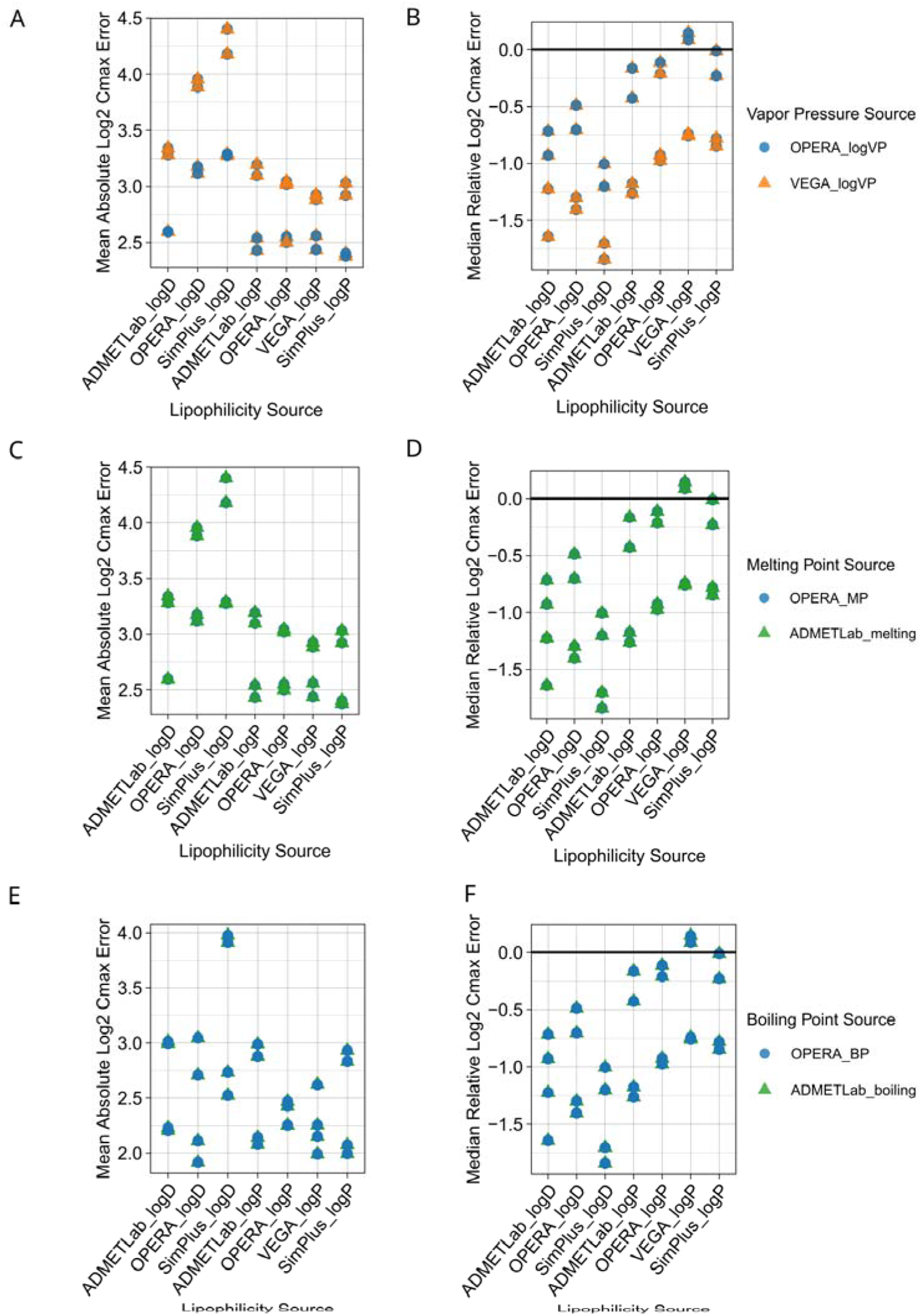
Comparison of systemic PK prediction performances for different vapour pressure, melting and boiling point prediction sources. Dermal absorption simulations of all parameterisation sources were examined against the collected *in vivo* dermal PK dataset. Solubility values were taken from SimPlus LogS. The first columns show Mean Absolute Log2 Cmax Errors (A, C, E) for different lipophilicity prediction sources. The second column shows Median Relative Log2 Cmax Errors (B, D, F). VP stands for vapour pressure, MP is melting point, BP is boiling point.

**Supplementary Fig. 2.**
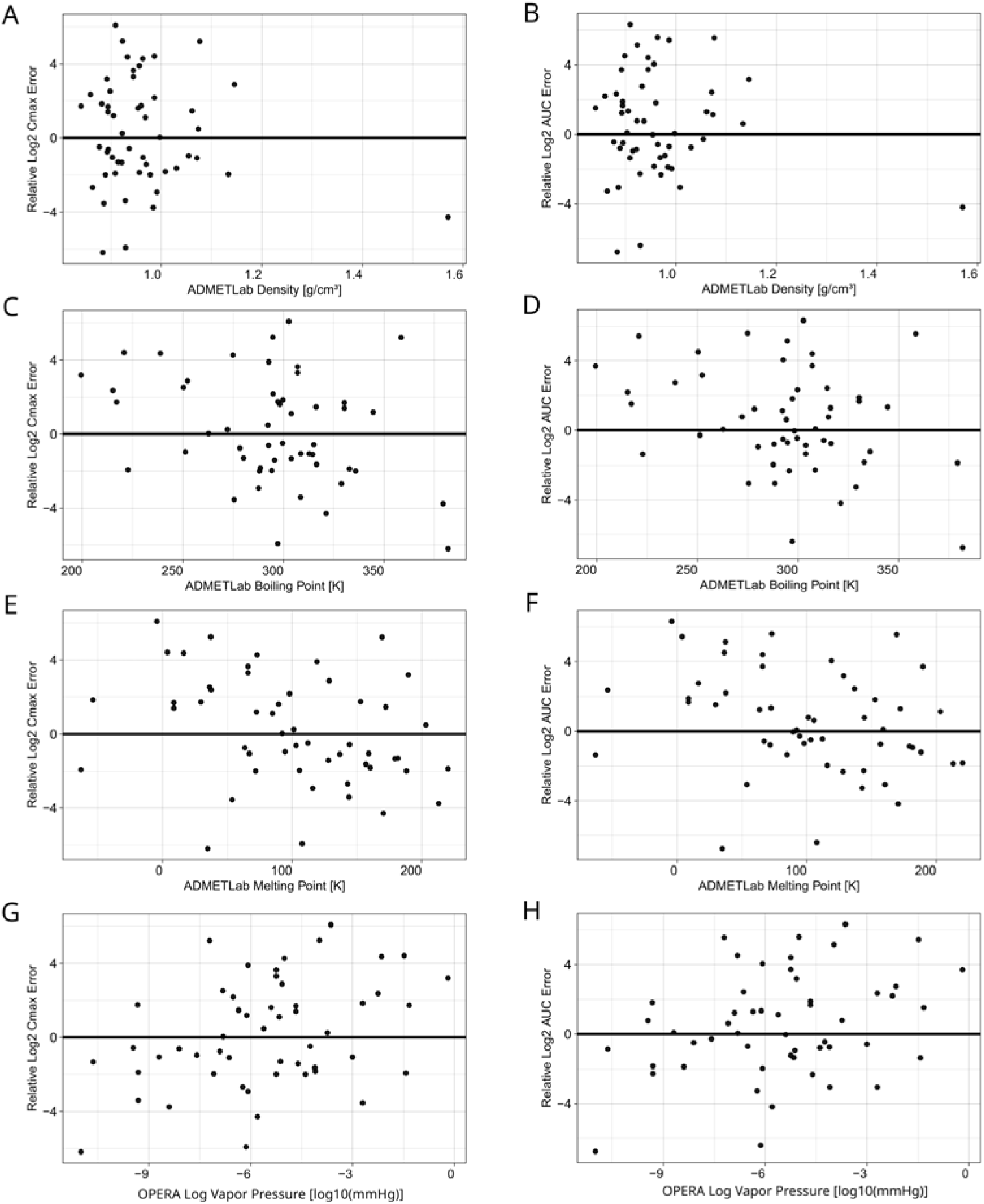
Correlation of PK prediction errors of the best dermal HT-PBK strategy with additional compound properties. The predicted compound parameters are compared against the Relative Log2 Cmax and AUC errors.

**Supplementary Fig. 3.**
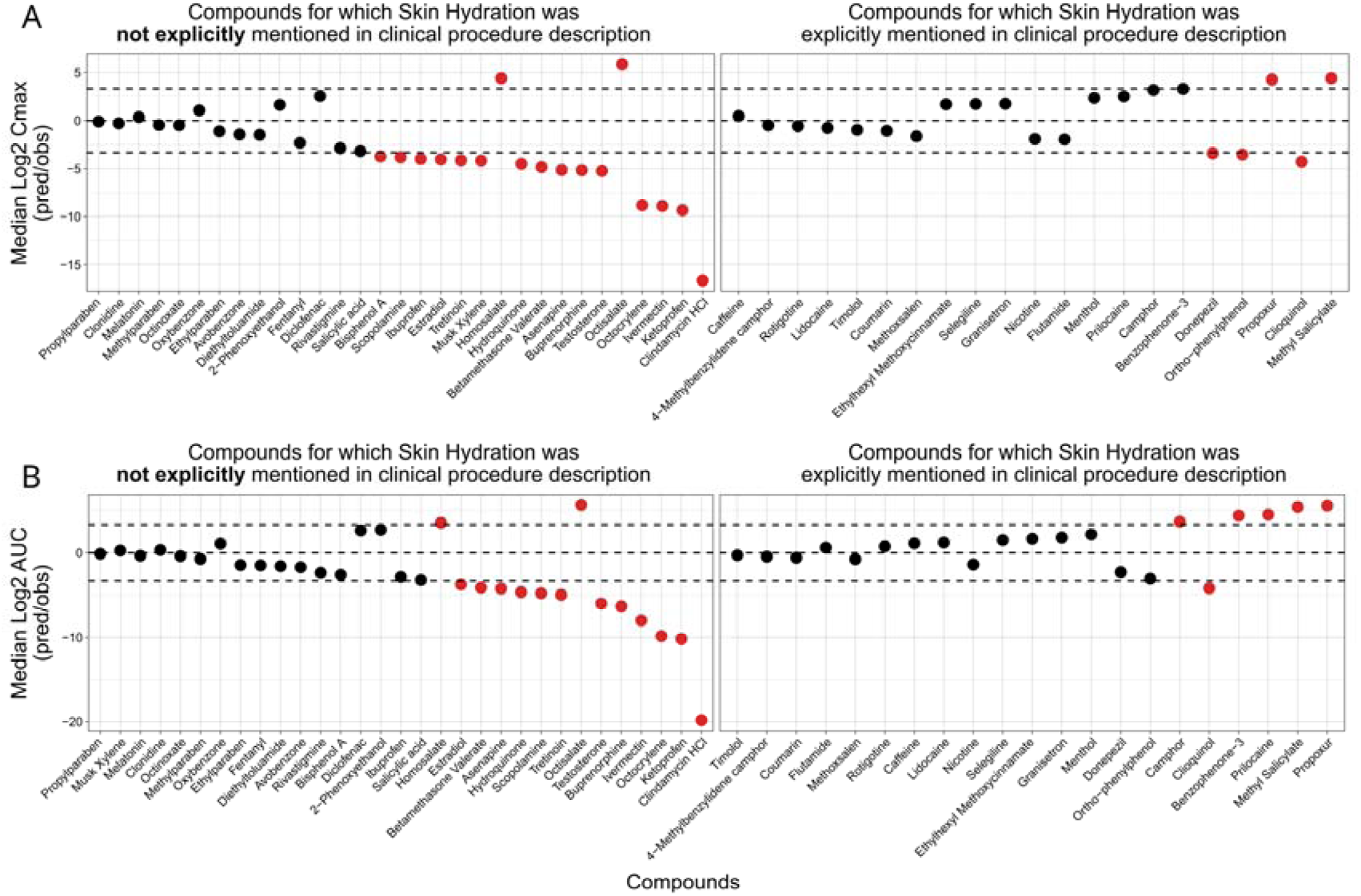
The effect of unspecified skin hydration status on systemic PK predictions. Many dermal PK studies did not explicitly mention whether application was performed on hydrated or non-hydrated skin. Shown results are for the assumption that lack of explicit mentioning relates to absence of skin hydration in study procedures (A) The Median Log2 Cmax predicted/observed values of each compound in the dataset. (B) The Median Log2 AUC predicted/observed values of each compound in the dataset. Plots are also split according to which information about the hydration status of the stratum corneum were provided in the clinical study information.

**Supplementary Fig. 4.**
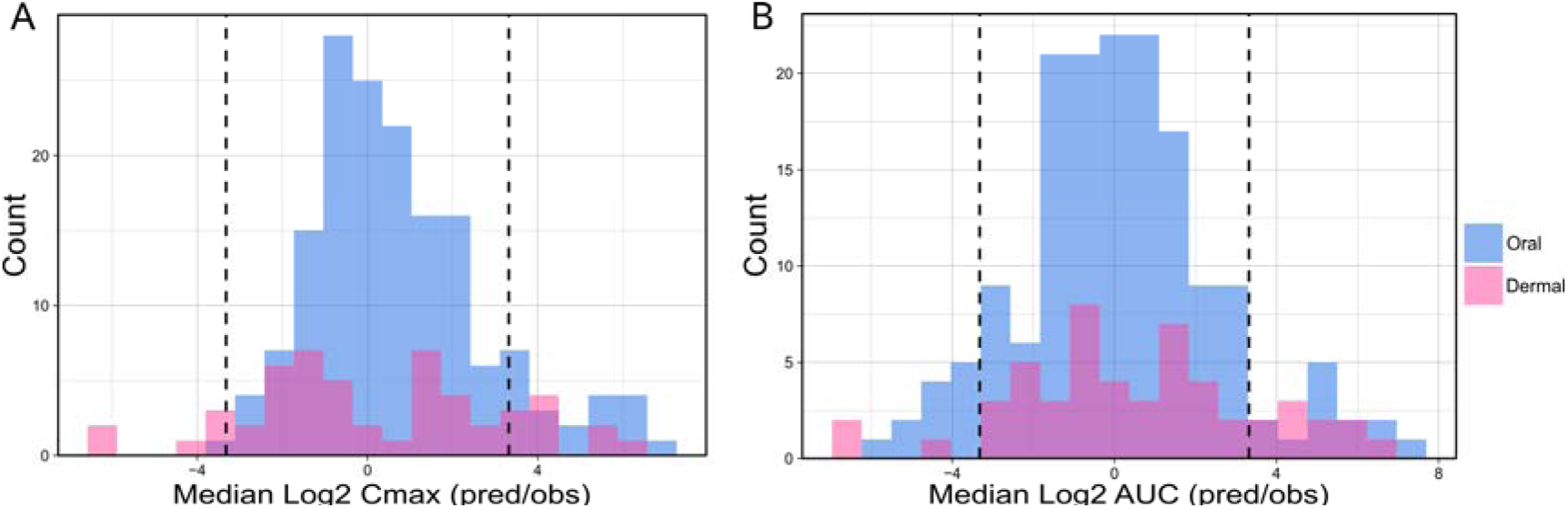
Comparison of non-normalised Cmax and AUC error distributions for dermal and oral PK predictions. (A) The Median Log2 predicted/observed Cmax values of dermally administered compounds (pink) were compared against orally administered compounds (blue). (B) The Median Log2 predicted/observed AUC values of dermally administered compounds (pink) were compared against orally administered compounds (blue). Oral HT-PBK prediction errors were taken from Geci et al. 2024.

## Notes

### Competing Interest Statement

The authors have declared no competing interest.

## References

1. Akomeah FK, Martin GP, Brown MB (2007) Variability in human skin permeability in vitro: comparing penetrants with different physicochemical properties. Journal of pharmaceutical sciences 96(4):824–834. doi: 10.1002/jps.20773

2. Anissimov YG, Roberts MS (2009) Diffusion modelling of percutaneous absorption kinetics: 4. Effects of a slow equilibration process within stratum corneum on absorption and desorption kinetics. Journal of pharmaceutical sciences 98(2):772–781. doi: 10.1002/jps.21461

3. Arora S, Clarke J, Tsakalozou E, Ghosh P, Alam K, Grice JE, Roberts MS, Jamei M, Polak S (2022) Mechanistic Modeling of In Vitro Skin Permeation and Extrapolation to In Vivo for Topically Applied Metronidazole Drug Products Using a Physiologically Based Pharmacokinetic Model. Molecular pharmaceutics 19(9):3139–3152. doi: 10.1021/acs.molpharmaceut.2c00229

4. Ball N, Bars R, Botham PA, Cuciureanu A, Cronin MTD, Doe JE, Dudzina T, Gant TW, Leist M, van Ravenzwaay B (2022) A framework for chemical safety assessment incorporating new approach methodologies within REACH. Arch Toxicol 96(3):743–766. doi: 10.1007/s00204-021-03215-9

5. Basak SC, Mills D, Mumtaz MM (2007) A quantitative structure-activity relationship (QSAR) study of dermal absorption using theoretical molecular descriptors. SAR and QSAR in Environmental Research 18(1-2):45–55. doi: 10.1080/10629360601033671

6. Benfenati E, Manganaro A, Gini G (2013) VEGA-QSAR: AI inside a platform for predictive toxicology. Popularize Artificial Intelligence 2013: Proceedings of the Workshop on Popularize Artificial Intelligence (PAI 2013)

7. Clarke JF, Thakur K, Polak S (2022) A mechanistic physiologically based model to assess the effect of study design and modified physiology on formulation safe space for virtual bioequivalence of dermatological drug products. Frontiers in pharmacology 13:1007496. doi: 10.3389/fphar.2022.1007496

8. Cross SE, Magnusson BM, Winckle G, Anissimov Y, Roberts MS (2003) Determination of the effect of lipophilicity on the in vitro permeability and tissue reservoir characteristics of topically applied solutes in human skin layers. The Journal of Investigative Dermatology 120(5):759–764. doi: 10.1046/j.1523-1747.2003.12131.x

9. Dancik Y, Miller MA, Jaworska J, Kasting GB (2013) Design and performance of a spreadsheet-based model for estimating bioavailability of chemicals from dermal exposure. Advanced drug delivery reviews 65(2):221–236. doi: 10.1016/j.addr.2012.01.006

10. Dancik Y, Zhang Y, Telaprolu KC, Polak S (2025) Physiologically based pharmacokinetic modelling of in vitro skin permeation of sunscreen actives under various experimental conditions. International Journal of Pharmaceutics 682:125977. doi: 10.1016/j.ijpharm.2025.125977

11. Denney WS, Duvvuri S, Buckeridge C (2015) Simple, Automatic Noncompartmental Analysis: The PKNCA R Package. Journal of pharmacokinetics and pharmacodynamics 42(1):11–107,S65. doi: 10.1007/s10928-015-9432-2

12. Dent MP, Vaillancourt E, Thomas RS, Carmichael PL, Ouedraogo G, Kojima H, Barroso J, Ansell J, Barton-Maclaren TS, Bennekou SH, Boekelheide K, Ezendam J, Field J, Fitzpatrick S, Hatao M, Kreiling R, Lorencini M, Mahony C, Montemayor B, Mazaro-Costa R, Oliveira J, Rogiers V, Smegal D, Taalman R, Tokura Y, Verma R, Willett C, Yang C (2021) Paving the way for application of next generation risk assessment to safety decision-making for cosmetic ingredients. Regulatory toxicology and pharmacology : RTP 125:105026. doi: 10.1016/j.yrtph.2021.105026

13. Egawa M, Hirao T, Takahashi M (2007) In vivo estimation of stratum corneum thickness from water concentration profiles obtained with Raman spectroscopy. Acta Dermato-Venereologica 87(1):4–8. doi: 10.2340/00015555-0183

14. Fu L, Shi S, Yi J, Wang N, He Y, Wu Z, Peng J, Deng Y, Wang W, Wu C, Lyu A, Zeng X, Zhao W, Hou T, Cao D (2024) ADMETlab 3.0: an updated comprehensive online ADMET prediction platform enhanced with broader coverage, improved performance, API functionality and decision support. Nucleic acids research 52(W1):W422–W431. doi: 10.1093/nar/gkae236

15. Geci R, Gadaleta D, Lomana MG de, Ortega-Vallbona R, Colombo E, Serrano-Candelas E, Paini A, Kuepfer L, Schaller S (2024) Systematic evaluation of high-throughput PBK modelling strategies for the prediction of intravenous and oral pharmacokinetics in humans. Arch Toxicol 98(8):2659–2676. doi: 10.1007/s00204-024-03764-9

16. Hamadeh A, Edginton A (2023) Efficient large-scale mechanism-based computation of skin permeability. Computational Toxicology 26:100263. doi: 10.1016/j.comtox.2023.100263

17. Hamadeh A, Nash JF, Bialk H, Styczynski P, Troutman J, Edginton A (2024) Mechanistic Skin Modeling of Plasma Concentrations of Sunscreen Active Ingredients Following Facial Application. Journal of pharmaceutical sciences 113(3):806–825. doi: 10.1016/j.xphs.2023.09.017

18. Hamadeh A, Troutman J, Najjar A, Edginton A (2022) A Mechanistic Bayesian Inferential Workflow for Estimation of In Vivo Skin Permeation from In Vitro Measurements. Journal of pharmaceutical sciences 111(3):838–851. doi: 10.1016/j.xphs.2021.11.028

19. Han P, Li X, Yang J, Zhang Y, Chen J (2024) Advancing Toxicity Predictions: A Review on in Vitro to in Vivo Extrapolation in Next-Generation Risk Assessment. Environment & health (Washington, D.C.) 2(7):499–513. doi: 10.1021/envhealth.4c00043

20. Jeong KM, Seo JY, Kim A, Kim YC, Baek YS, Oh CH, Jeon J (2023) Ultrasonographic analysis of facial skin thickness in relation to age, site, sex, and body mass index. Skin Research and Technology 29(8):e13426. doi: 10.1111/srt.13426

21. Kasting GB, Miller MA (2006) Kinetics of finite dose absorption through skin 2: volatile compounds. Journal of pharmaceutical sciences 95(2):268–280. doi: 10.1002/jps.20497

22. Kuepfer L, Niederalt C, Wendl T, Schlender J-F, Willmann S, Lippert J, Block M, Eissing T, Teutonico D (2016) Applied Concepts in PBPK Modeling: How to Build a PBPK/PD Model. CPT: pharmacometrics & systems pharmacology 5(10):516–531. doi: 10.1002/psp4.12134

23. Li H, Reynolds J, Sorrell I, Sheffield D, Pendlington R, Cubberley R, Nicol B (2022) PBK modelling of topical application and characterisation of the uncertainty of Cmax estimate: A case study approach. Toxicology and Applied Pharmacology 442:115992. doi: 10.1016/j.taap.2022.115992

24. Mansouri K, Grulke CM, Judson RS, Williams AJ (2018) OPERA models for predicting physicochemical properties and environmental fate endpoints. Journal of Cheminformatics 10(1):10. doi: 10.1186/s13321-018-0263-1

25. Najjar A, Schepky A, Krueger C-T, Dent M, Cable S, Li H, Grégoire S, Roussel L, Noel-Voisin A, Hewitt NJ, Cardamone E (2021) Use of Physiologically-Based Kinetics Modelling to Reliably Predict Internal Concentrations of the UV Filter, Homosalate, After Repeated Oral and Topical Application. Frontiers in pharmacology 12:802514. doi: 10.3389/fphar.2021.802514

26. Narita I, Todo H, Fujiwara C, Teramae H, Oshizaka T, Itakura S, Komatsu S, Takayama K, Sugibayashi K (2025) In silico model to predict dermal absorption of chemicals in finite dose conditions. The Journal of Toxicological Sciences 50(4):171–186. doi: 10.2131/jts.50.171

27. OECD (2021) Guidance document on the characterisation, validation and reporting of Physiologically Based Kinetic (PBK) models for regulatory purposes. OECD Series on Testing and Assessment No. 331, Paris

28. Oltulu P, Ince B, Kokbudak N, Findik S, Kilinc F (2018) Measurement of epidermis, dermis, and total skin thicknesses from six different body regions with a new ethical histometric technique. Turkish Journal of Plastic Surgery 26(2):56. doi: 10.4103/tjps.TJPS_2_17

29. Paini A, Leonard JA, Joossens E, Bessems JGM, Desalegn A, Dorne JL, Gosling JP, Heringa MB, Klaric M, Kliment T, Kramer NI, Loizou G, Louisse J, Lumen A, Madden JC, Patterson EA, Proença S, Punt A, Setzer RW, Suciu N, Troutman J, Yoon M, Worth A, Tan YM (2019) Next generation physiologically based kinetic (NG-PBK) models in support of regulatory decision making. Computational toxicology (Amsterdam, Netherlands) 9:61–72. doi: 10.1016/j.comtox.2018.11.002

30. Paini A, Tan Y-M, Sachana M, Worth A (2021) Gaining acceptance in next generation PBK modelling approaches for regulatory assessments - An OECD international effort. Computational toxicology (Amsterdam, Netherlands) 18:100163. doi: 10.1016/j.comtox.2021.100163

31. Patel N, Clarke JF, Salem F, Abdulla T, Martins F, Arora S, Tsakalozou E, Hodgkinson A, Arjmandi-Tash O, Cristea S, Ghosh P, Alam K, Raney SG, Jamei M, Polak S (2022) Multi-phase multi-layer mechanistic dermal absorption (MPML MechDermA) model to predict local and systemic exposure of drug products applied on skin. CPT: pharmacometrics & systems pharmacology 11(8):1060–1084. doi: 10.1002/psp4.12814

32. Poet TS, McDougal JN (2002) Skin absorption and human risk assessment. Chemico-biological interactions 140(1):19–34. doi: 10.1016/s0009-2797(02)00013-3

33. Punt A, Baltazar MT, Nicol B, Cable S, Hewitt NJ, Cubberley R, Spriggs S, Dent MP, Li H (2025) Building confidence in PBK model predictions in the absence of human kinetic data: Benzophenone-4 case study. ALTEX. doi: 10.14573/altex.2501211

34. Punt A, Bouwmeester H, Blaauboer BJ, Coecke S, Hakkert B, Hendriks DFG, Jennings P, Kramer NI, Neuhoff S, Masereeuw R, Paini A, Peijnenburg AACM, Rooseboom M, Shuler ML, Sorrell I, Spee B, Strikwold M, van der Meer AD, van der Zande M, Vinken M, Yang H, Bos PMJ, Heringa MB (2020) New approach methodologies (NAMs) for human-relevant biokinetics predictions. Meeting the paradigm shift in toxicology towards an animal-free chemical risk assessment. ALTEX 37(4):607–622. doi: 10.14573/altex.2003242

35. R Core Team (2024) R: a language and environment for statistical computing. https://www.R-project.org/

36. Roberts MS, Cross SE, Anissimov YG (2004) Factors affecting the formation of a skin reservoir for topically applied solutes. Skin Pharmacology and Physiology 17(1):3–16. doi: 10.1159/000074057

37. Rohatgi A (2024) Webplotdigitizer: Version 5.2

38. Sandby-Møller J, Poulsen T, Wulf HC (2003) Epidermal thickness at different body sites: relationship to age, gender, pigmentation, blood content, skin type and smoking habits. Acta Dermato-Venereologica 83(6):410–413. doi: 10.1080/00015550310015419

39. Taylor K, Rego Alvarez L (2020) Regulatory drivers in the last 20 years towards the use of in silico techniques as replacements to animal testing for cosmetic-related substances. Computational Toxicology 13:100112. doi: 10.1016/j.comtox.2019.100112

40. van Mulder T, Koeijer M de, Theeten H, Willems D, van Damme P, Demolder M, Meyer G de, Beyers K, Vankerckhoven V (2017) High frequency ultrasound to assess skin thickness in healthy adults. Vaccine 35(14):1810–1815. doi: 10.1016/j.vaccine.2016.07.039

41. Wetmore BA, Wambaugh JF, Ferguson SS, Sochaski MA, Rotroff DM, Freeman K, Clewell HJ, Dix DJ, Andersen ME, Houck KA, Allen B, Judson RS, Singh R, Kavlock RJ, Richard AM, Thomas RS (2012) Integration of dosimetry, exposure, and high-throughput screening data in chemical toxicity assessment. Toxicological sciences : an official journal of the Society of Toxicology 125(1):157–174. doi: 10.1093/toxsci/kfr254

42. Willmann S, Lippert J, Sevestre M, Solodenko J, Fois F, Schmitt W (2003) PK-Sim®: a physiologically based pharmacokinetic ‘whole-body’ model. BIOSILICO 1(4):121–124. doi: 10.1016/S1478-5382(03)02342-4

43. Y. L, K. H (2002) Skin thickness of Korean adults. Surgical and Radiologic Anatomy 24(3-4):183–189. doi: 10.1007/s00276-002-0034-5

